# *In Silico* study of clinical implication of markers associated with PTHrP regulatory mechanisms and linked to angiogenesis and EMT program of colorectal cancer

**DOI:** 10.64898/2026.04.15.718767

**Authors:** Pedro Carriere, María Belén Novoa Díaz, Cintia Birkenstok, Claudia Gentili

## Abstract

Parathyroid hormone-related peptide (PTHrP), encoded by PTHLH, has been implicated in tumor progression through its involvement in epithelial-mesenchymal transition (EMT), angiogenesis, and tumor cell migration. Previous experimental studies suggest that PTHrP may promote these processes in colorectal cancer (CRC), partly through the modulation of factors such as secreted protein acidic and rich in cysteine (SPARC) and vascular endothelial growth factor (VEGFA). These events play a key role in the acquisition of an aggressive phenotype in our experimental models. In this study, we performed an integrative *in silico* analysis of multiple transcriptomic datasets to investigate the potential role of PTHLH in CRC. Differential expression analysis identified a set of consistently dysregulated genes across independent datasets. Functional enrichment and network analyses revealed that PTHLH expression is associated with biological processes related to extracellular matrix remodeling, EMT, and angiogenesis. Correlation analyses showed a positive association between PTHLH and SPARC expression, while network-based approaches suggested a potential functional connection with VEGFA. To assess the clinical relevance of these findings, survival analysis was performed using publicly available datasets. High expression levels of PTHLH, SPARC, and VEGFA were significantly associated with reduced overall survival in patients. Notably, a combined gene signature based on these three factors demonstrated a stronger prognostic effect than individual genes, indicating enhanced predictive value. These findings suggest that PTHrP is associated with molecular pathways involved in tumor progression and, together with SPARC and VEGF, may contribute to a coordinated regulatory axis with prognostic relevance in CRC, warranting further experimental validation.

## 1. Introduction

According to statistics from the Global Cancer Observatory (Globocan), colorectal cancer (CRC) is the third most common cancer after breast and lung cancer, accounting for 12.1% of all tumors in both sexes [1]. Because it is a molecularly heterogeneous pathology, in 2015 an International Committee reported a new classification of these tumors into four consensus molecular subtypes (CMS) that present distinctive features. The CMS1 (Immune, 14% of cases) is a group with microsatellite instability and strong immune activation; the CMS2 (Canonical, 37% of cases) have epithelial features, chromosome instability and marked activation of Wnt (Wingless-related integration site) and MYC (MYC Proto-Oncogene, BHLH Transcription Factor) signaling; CMS3 (Metabolic, 13% of cases) have epithelial attributes with evident metabolic dysregulation; and the CMS4 (Mesenchymal, 23% of cases) have prominent activation of transforming growth factor β (TGF-β), stromal invasion and angiogenesis [2]. In particular, CMS4-type cancers, often diagnosed at advanced stages (III and IV), have a poor prognosis with worse 5-year overall survival (62% of patients) and relapse-free survival (60% of patients) compared to the other molecular subtypes [2]. This CRC classification highlights the implication of the tumor microenvironment (TME) and raises the need to understand the factors involved in its progression. Among them, parathyroid hormone-related peptide (PTHrP), a cytokine present in the TME, has been related to the initiation and progression of different tumors [3-5].

The results obtained by our group reveal that PTHrP, on intestinal tumor cells promotes survival, cell cycle progression, proliferation, migration, cancer stem cell (CSC) features and resistance to drugs commonly used in CRC treatment [3,6-9]. Furthermore, we found that this peptide increases the expression of the pro-angiogenic factors VEGF (Vascular endothelial growth factor), HIF-1α (Hypoxia-inducible factor 1 subunit alpha), MMP-7 (Matrix metallopeptidase 7) and MMP-9 (Matrix metallopeptidase 9) in CRC-derived cell lines Caco-2 and HCT116 and promotes tumor-associated angiogenesis through VEGF. These effects are mediated by the activation of PI3K/Akt and ERK1/2 pathways [10]. Regarding the epithelial-mesenchymal transition (EMT) phenomenon, in HCT116 cells, we observed that PTHrP induces morphological changes from an epithelial to a mesenchymal phenotype. This event was accompanied by a decrease in the expression of the epithelial markers CK-18 (Cytokeratin 18) and E-cadherin, as well as an increase in the expression of N-cadherin and ZEB-1 (Zinc finger E-box-binding homeobox 1) proteins associated with a mesenchymal state [11]. Besides, conditioned medium from PTHrP-treated HMEC-1 endothelial cells (CME) decreases E-cadherin expression in HCT116 cells. Also, the CME favors a mesenchymal phenotype, suggesting that factors present in this medium could participate in the EMT program in CRC cells. We observed that PTHrP modulates the expression of SPARC (secreted protein acidic and rich in cysteine), a protein that participates in extracellular matrix remodeling and that is associated with the CRC aggressive phenotype in HCT116 cells [11]. In HMEC-1 endothelial cells, the treatment with PTHrP also increases the expression of this protein and its secretion into the extracellular medium. Furthermore, in CRC cells, exogenous SPARC exerts a synergistic effect with PTHrP on cell migration and molecular events related to EMT such as the decrease in E-cadherin protein expression. Finally, the peptide administered to an *in vivo* model promotes morphological and molecular changes that validate these events observed *in vitro* [11].

Based on the results obtained *in vitro* and *in vivo*, this work aims to determine the relevance in the clinical context of markers associated with the regulatory mechanisms of PTHrP in CRC models, through an in silico analysis of data from samples of CRC patients published on online platforms.

## 2. Materials and methods

### 2.1 Microarray gene expression data

The GEO (Gene Expression Omnibus) platform was used to obtain four microarray expression data sets from CRC patients. Data were obtained from the following accessions: GSE41258: gene expression platform Affymetrix Human Genome U133A Array - 54 normal samples and 186 primary tumor samples [12]; GSE37364: gene expression platform Affymetrix Human Genome U133 Plus 2.0 Array - 38 normal samples and 27 primary tumor samples; GSE68468: gene expression platform Affymetrix Human Genome U133A Array - 53 normal samples and 186 primary tumor samples [13], and GSE9348: gene expression platform Affymetrix Human Genome U133 Plus 2.0 Array - 12 normal samples and 70 primary tumor samples [14]. From each dataset, all primary tumor and normal samples were selected (accessed on 20 April 2022).

### 2.2 Obtaining and identifying differentially expressed genes (DEGs)

Differential expression analysis was performed with R software using the fold change (FC) criterion. To obtain the differentially expressed genes (DEGs), a threshold of FC>2 (an expression level greater than twofold) corresponding to a Log_2_FC>1 was selected for the positively regulated genes. For negatively regulated genes, a threshold of FC<1/2 (less than half an expression level) was selected, corresponding to a Log_2_FC<-1.

### 2.3 Enrichment Analysis and Expression Platforms Consulting

The obtained DEGs were incorporated into the FunRich 3.1.3 program (“Functional Enrichment Analysis Tool”) [15] and analyzed with functional enrichment using Enrichr [16]. To explore the clinical implication and interaction between the evaluated genes, we explored the EMTome (http://www.emtome.org/), GEPIA2 (“Gene Expression Profiling Interactive Analysis 2”), http://gepia2.cancer-pku.cn/#index [17] and STRING 11.5, https://string-db.org/ [18] databases. (accessed on 5 May 2022).

### 2.4 Interaction network construction and topological analysis

Cytoscape 3.8.2 and its stringApp [19] were used to integrate the expression data of the DEGs into networks imported from the STRING protein-protein interaction (PPI) database to visualize central molecular networks related to CRC. The detection of hub genes of the network was determined using the CytoNCA app from Cytoscape, using the criteria of Degree, Closeness, Betweenness and Eigenvector [17]. Clusters of highly interconnected nodes from this network were obtained using the Cytoscape plugin MCODE (molecular complex detection algorithm) [20] (accessed on 5 May 2022).

### 2.5 Survival analysis

Parathyroid hormone-related peptide gene (PTHLH) (210355_at), SPARC (212667_at), and VEGFA (210512_s_at) gene expression were evaluated for their prognostic significance in CRC patients using the Kaplan–Meier Plotter (https://kmplot.com/analysis/) [21]. Data were derived from multiple independent cohorts integrated within the platform. For each gene, patients were stratified into high- and low-expression groups. To ensure robustness, survival analyses were performed using both the auto-selected best cutoff and the median expression threshold. Overall survival (OS) was used as the primary endpoint. Survival curves were generated using the Kaplan-Meier method, and statistical significance between groups was assessed using the log-rank test. Hazard ratios (HRs) with 95% confidence intervals were calculated using a Cox proportional hazards model. To evaluate the combined prognostic value of these genes, a multigene signature was constructed using the mean expression of PTHLH, SPARC, and VEGFA, as implemented in the Kaplan-Meier Plotter platform. To ensure robustness and reduce potential bias associated with optimal cutoff selection, survival analyses were additionally evaluated using the median expression threshold.

### 2.6 Statistical analysis

Comparisons of expressions from microarray data were analyzed by moderated t-student with a probability of p<0.05 and a probability adjusted by Benjamini and Hochberg or false discovery rate (FDR) of p adj<0.05 [22]. In the clinical samples, the correlation between the gene expression of PTHrP (PTHLH), VEGFA (VEGFA), and SPARC was evaluated using the Pearson coefficient (R) and the Spearman test. These statistical analyses were performed using the R programming language version 4.1.1. (Available at: https://www.R-project.org/). In all cases, a p-value < 0.05 was considered statistically significant.

## 3 Results

### 3.1 Identification of DEGs in CRC samples and their association with angiogenesis and EMT

Initially, the online database GEO and the R software were used to analyze four gene expression data sets of intestinal samples from patients with CRC compared to samples from healthy patients. As seen in **Figure 1A**, the analysis of tumor samples and normal colorectal samples resulted in the obtaining of DEGs in the four data sets. Then, the DEGs (Log_2_FC>1 and Benjamini and Hochberg adjusted p < 0.05) of each data set were incorporated into the FunRich 3.1.3 software to determine the genes recognized and shared by the four data sets (**Figure 1B** and **1C**). The list of positively and negatively regulated genes that were common to the four datasets is shown in **Table 1**. In particular, in datasets GSE37364 and GSE9348, the comparison analysis of normal and tumor samples revealed that PTHLH is possibility regulated in tumor samples (**Table 2**).

**Table 1.**
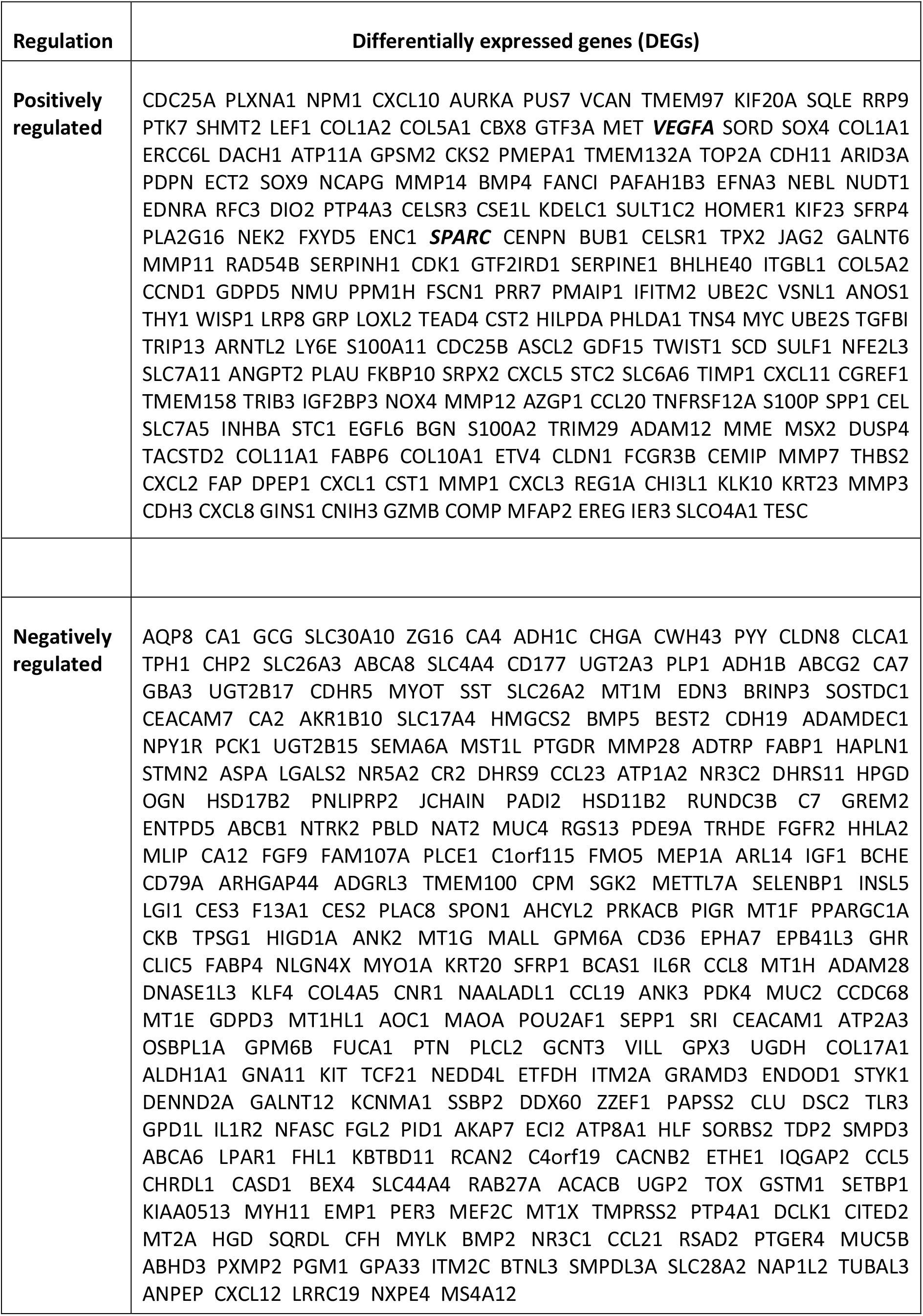
List of positively and negatively regulated genes common to the four CRC samples data sets.

**Table 2.** The PTHrP gene (PTHLH) is positively regulated in two of the four CRC samples data sets.

| Accession | p adjusted | p Value | t | log<sub>2</sub>FC | Gene |
| --- | --- | --- | --- | --- | --- |
| **GSE37364** | 5,76x10<sup>-8</sup> | 4,21x10<sup>-9</sup> | 67.323.871 | 2.34084 | PTHLH |
| **GSE9348** | 1,56x10<sup>-3</sup> | 2,45x10<sup>-4</sup> | 38.334.384 | 1.5279 | PTHLH |

**Figure 1.**
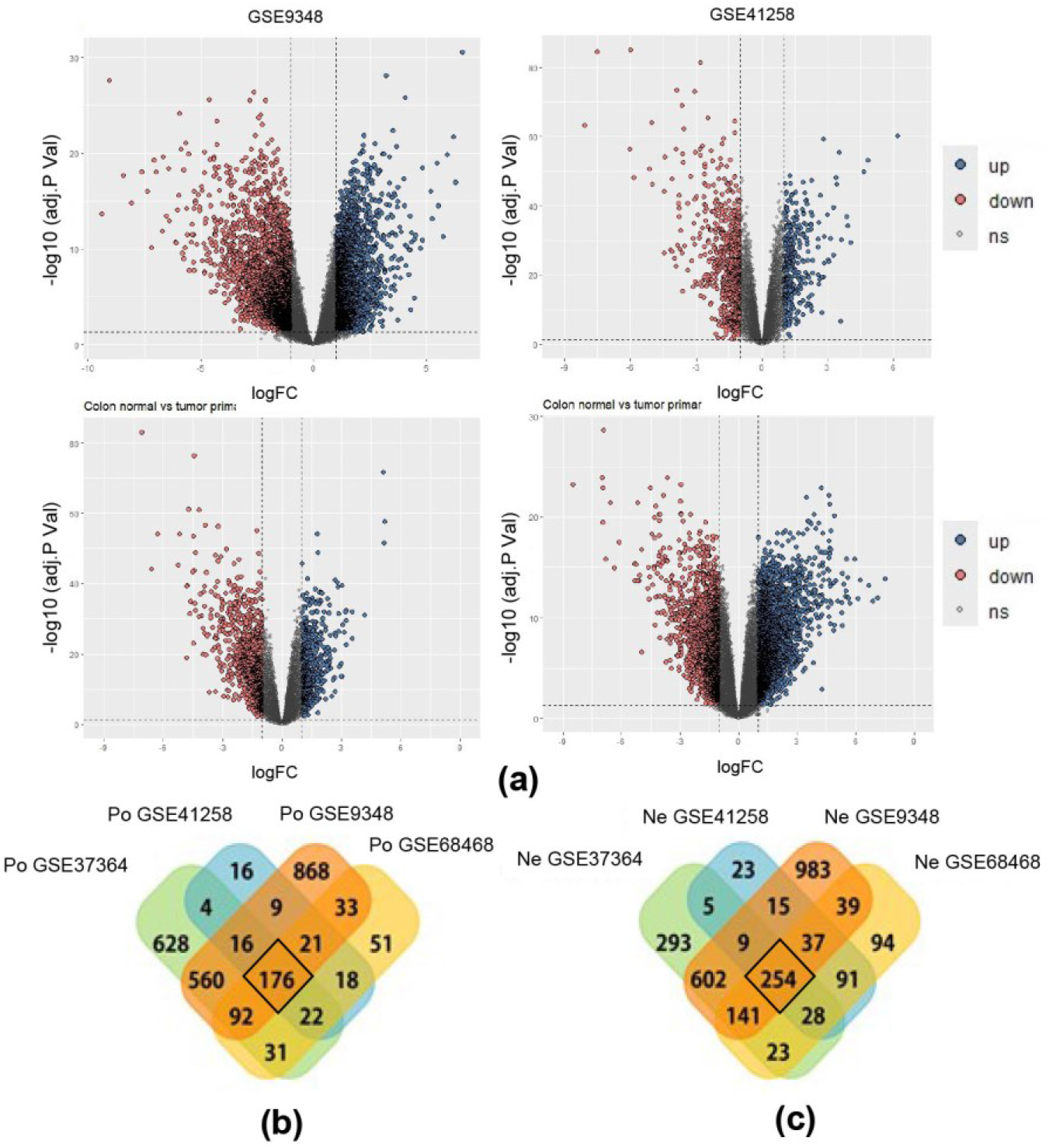
Four CRC patient datasets share differentially expressed genes in tumor samples. **(a)** Volcano diagram. DEGs were obtained in the tumor tissue of each dataset (blue: positively regulated; red: negatively regulated); **(b)** Venn diagram obtained employing FunRich software. 176 positively regulated (Po) and **(c)** 254 negatively regulated (Ne) genes were identified that were common to tumor samples from the four CRC datasets, with significant expression change (Log_2_FC>1; Log_2_FC<-1 and adjusted p < 0.05).

To gain a deeper understanding of the function of the DEGs shared by the four datasets, their participation in biological processes, the signaling pathways in which they are involved, and their subcellular localization were assessed. For this purpose, the DEGs were incorporated into the Enrichr platform [16], which includes different databases for functional enrichment. As seen in **Figure 2**, the positively regulated genes in CRC are significantly associated with extracellular matrix organization processes (p<2.11x10^-16^), EMT pathways (p<1.95x10^-28^), NF-kB (Nuclear factor kappa-light-chain-enhancer of activated B cells) signaling pathways (p<2.37x10^-12^), angiogenesis pathways (p<2.44x10^-8^) and KRAS (Kirsten rat sarcoma virus) overactivated signaling (p<2.16x10^-7^). Also, they are associated with VEGFA complexes (p<5.34x10^-25^), among others. On the other hand, the negatively regulated genes by all data sets showed an association with metal ion metabolism (p<5.35x10^-14^), pathways associated with KRAS signaling (7.88x10^-10^), fatty acid metabolism (8.68X10-7) and association with the extracellular region (p<6.23x10^-18^) and apical complexes (p<3.17x10^-17^) (**Figure 3**). These results highlight the importance of the pathways involved in EMT and angiogenesis in the genesis/progression of CRC.

**Figure 2.**
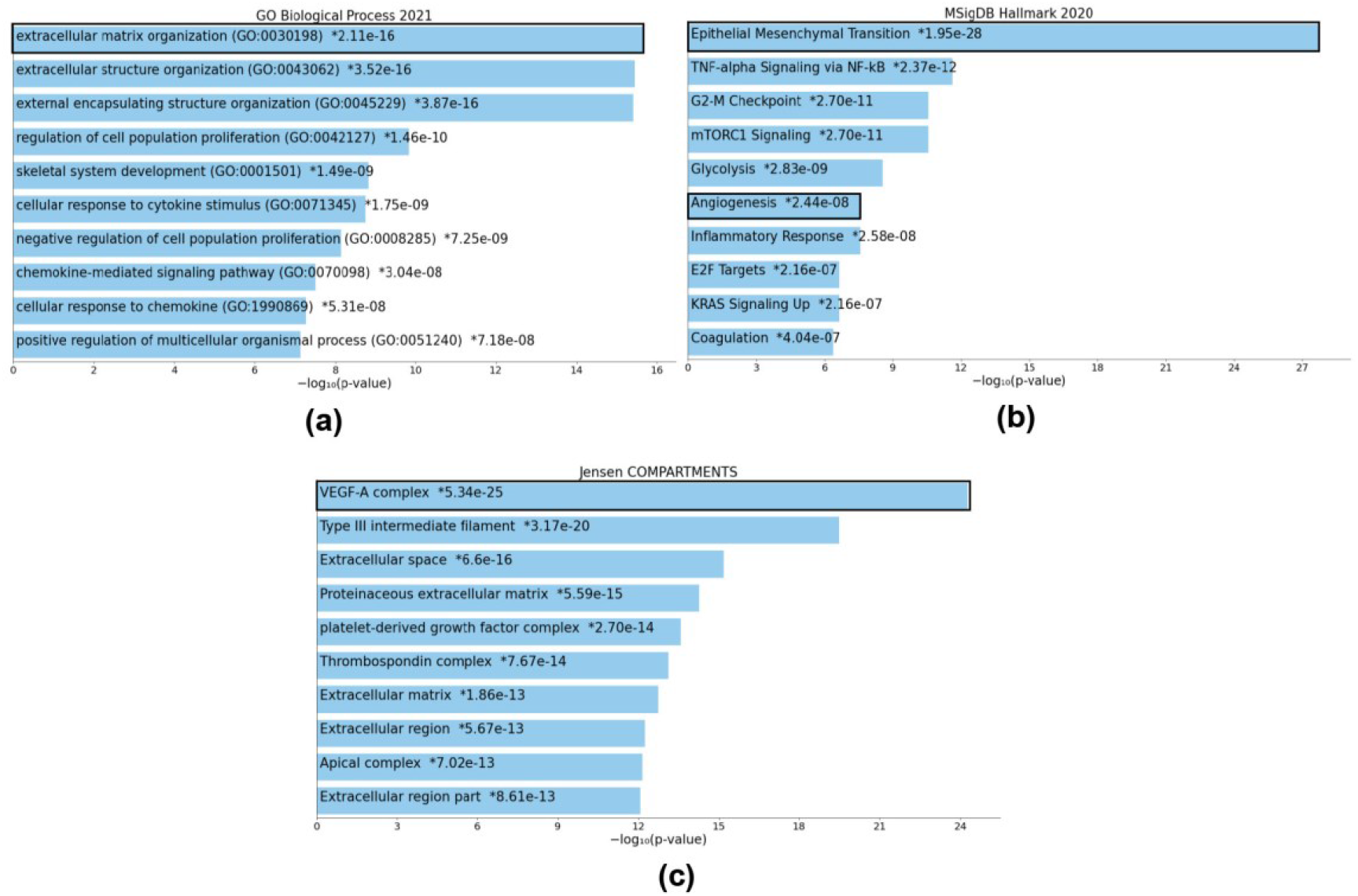
Positively regulated genes in CRC are enriched in pathways associated with angiogenesis and EMT. Using the Enrichr platform, the functional enrichment query was performed for **(a)** biological processes, **(b)** signaling pathways, and **(c)** subcellular compartments of the 176 positively regulated genes shared by the four CRC data sets. The bar graph shows the associated probability in each case expressed as – log_10_ (p-value).

**Figure 3.**
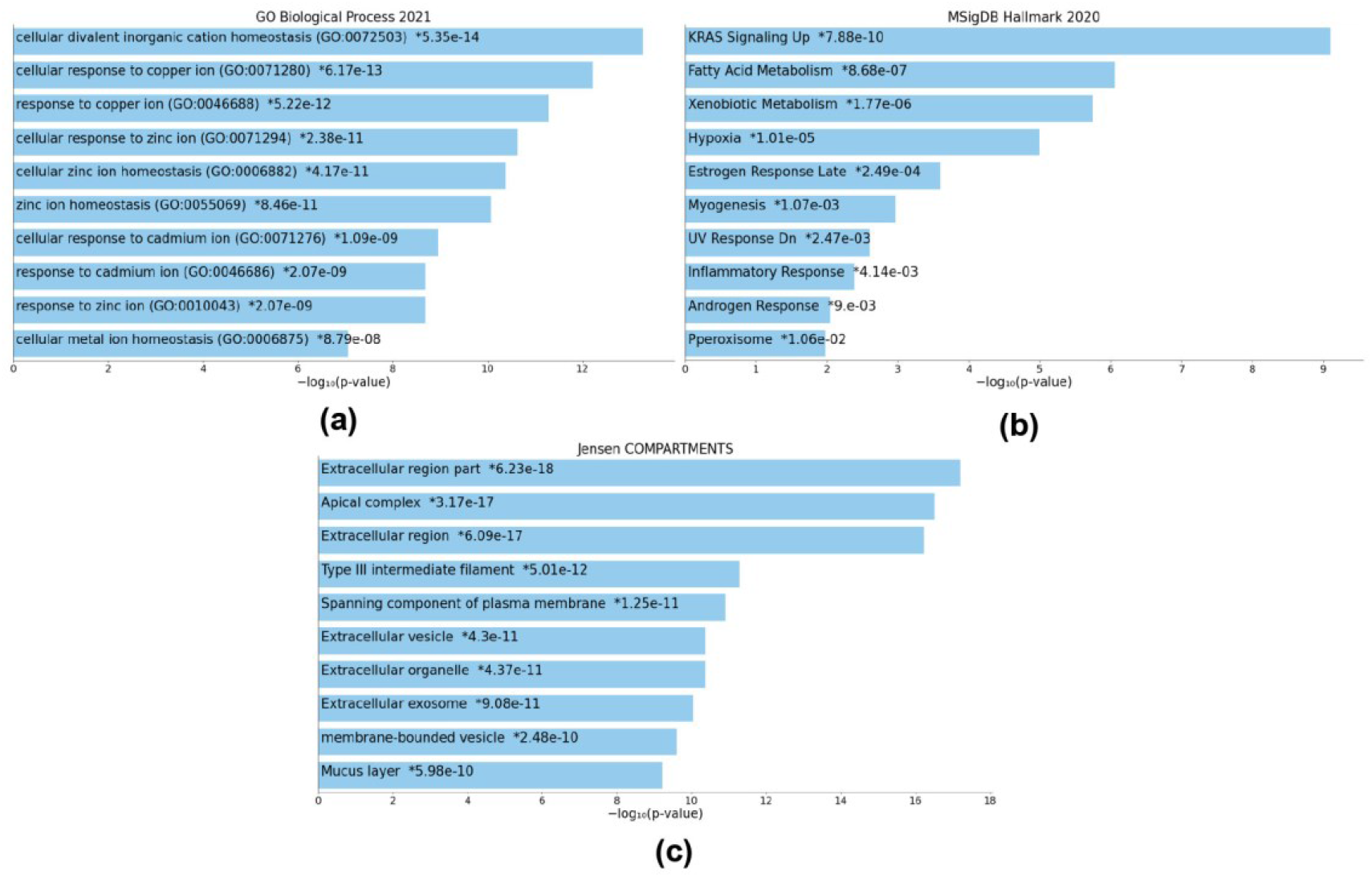
Negatively regulated genes in CRC are enriched in pathways associated with CRC progression not yet evaluated by our research group. Using the Enrichr platform, the functional enrichment query was performed by **(a)** biological processes, **(b)** signaling pathways, and **(c)** subcellular compartments, of the 254 negatively regulated genes shared by the four CRC data sets. The bar graph shows the associated probability in each case expressed as – log_10_ (p-value).

### 3.2 Correlation analysis of the PTHrP gene with angiogenesis and EMT

The relationship of PTHrP in different types of cancer and its association with the processes of EMT and angiogenesis was then explored accessing to the enormous amount of information currently available in different clinical sample analysis platforms. Initially, the EMTome platform [22] was used, which contains signatures of genes associated with EMT derived from the Cancer Cell Line Encyclopedia (CCLE) [24], from metastatic patient samples (MET500) [25], from clinical samples taken from the The Cancer Genome Atlas (TCGA) repository [26] and from data from the Human Protein Atlas (HPA) [27].

The gene PTHLH was entered into this platform as a search and different data classes were obtained. The top 10 correlation values of the PTHLH gene search are associated with EMT, angiogenesis, coagulation, apical functions, and KRAS signaling (**Figure 4**). By exploring the different cellular processes related to PTHLH expression, it was observed that in colon cancer (COAD) and rectal cancer (READ) this gene has a high positive correlation with pathways involved in EMT, angiogenesis, coagulation, apical junctions, TGF-β signaling and KRAS signaling, and other biological functions with lesser correlation (**Figure 5**).

**Figure 4.**
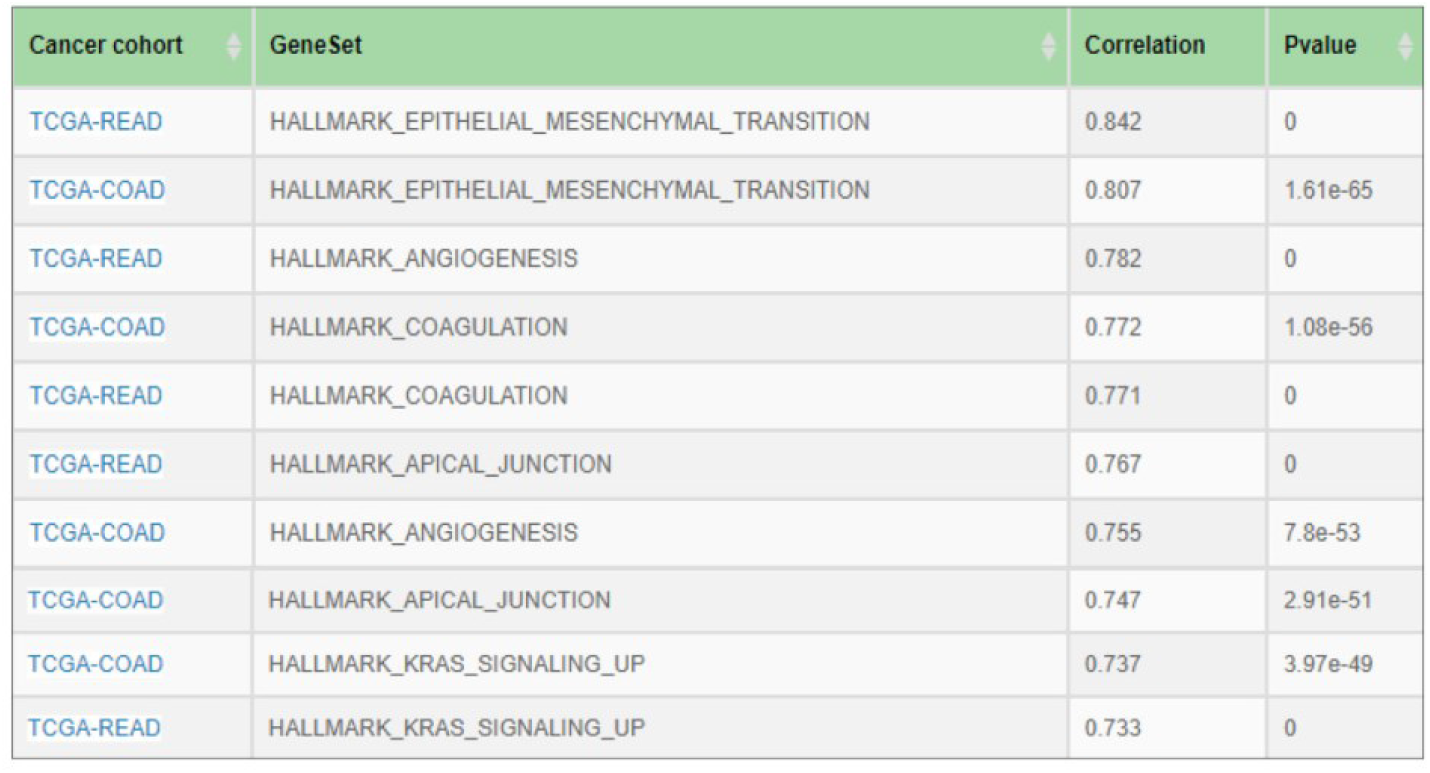
Core processes associated with the PTHrP gene (PTHLH) in the TCGA clinical sample repository.

**Figure 5.**
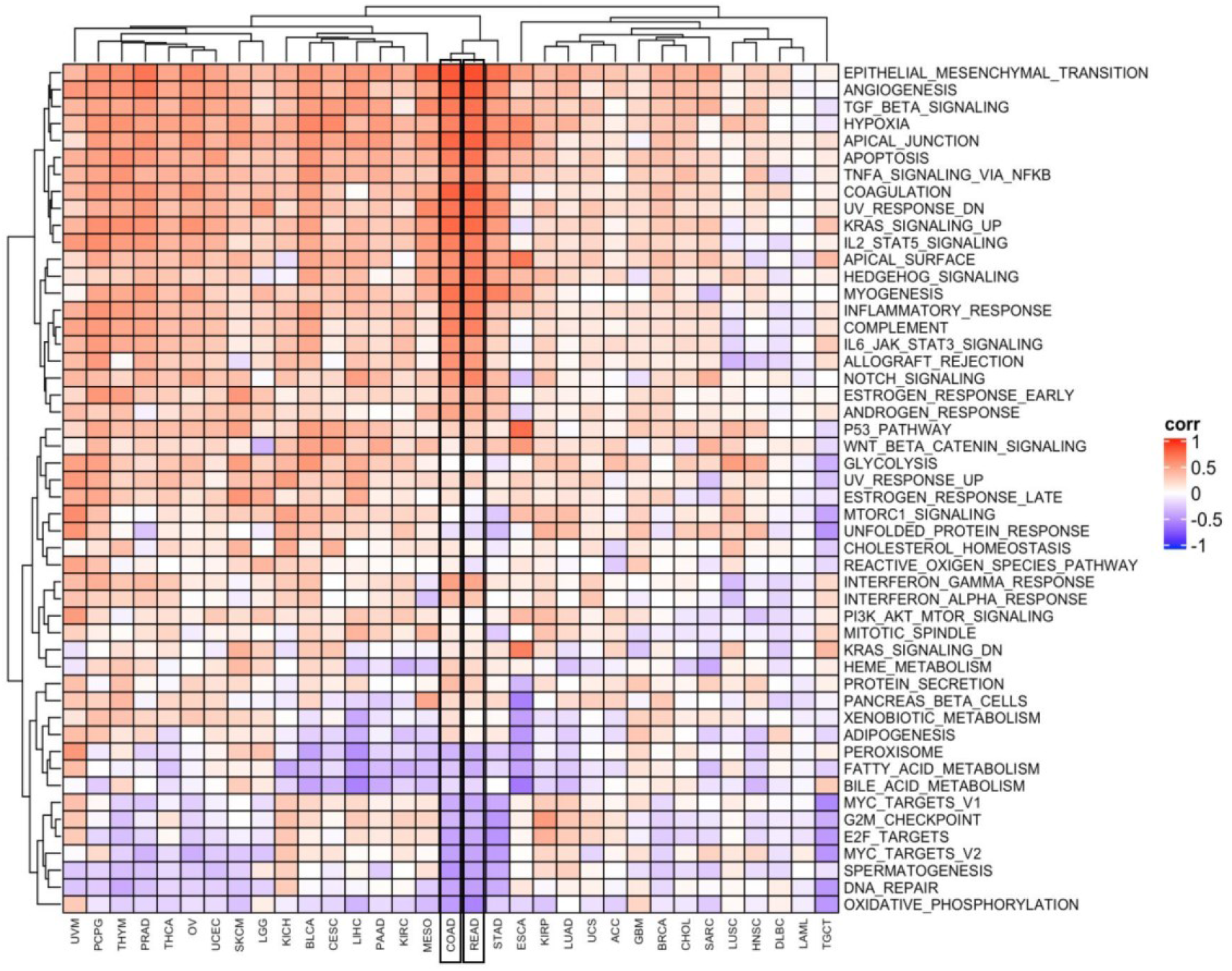
PTHLH gene is significantly correlated with EMT and angiogenesis in colon cancer (COAD) and rectal cancer (READ). The correlation diagram shows the correlation with different events in different tumor types (Hallmarks Geneset from EMTome).

To determine in clinical samples whether there is an association between PTHrP and the markers modulated by this cytokine in our experimental CRC models, an analysis was performed using the Pearson correlation test (R), using the online database GEPIA2 [17], which includes 9,736 tumor samples and 8,587 normal samples from the TCGA [26] and Genotype-Tissue Expression (GTEx) projects [28]. As seen in **Figure 6**, PTHLH gene expression is positively and significantly correlated with markers associated with angiogenesis, extracellular matrix remodeling, and cell migration such as HIF-1 gene expression (R=0.37; p=3.5x10^-10^), MMP-7 (R=0.29; p= 7.2x10-7) and MMP-9 (R=0.25; p=2.1x10^-5^) in colon cancer samples. Nevertheless, no significant correlation was found in the data analyzed regarding the expression of PTHLH and VEGFA in this database. On the other hand, a positive and significant correlation was also found between PTHLH and mesenchymal markers previously evaluated by our group [11], ZEB1 (R=0.35; p=3.9X10^-9^), N-cadherin (R=0.41; p=1.1x10^-12^) and SPARC (R=0.58; p=0), but not between PTHLH and the gene expression of epithelial markers such as E-cadherin (**Figure 7**). In addition, using the clinical samples analyzed where the overexpression of the PTHLH gene was found (GSE9348 and GSE37364), it was evaluated whether its overexpression correlates with the expression of VEGF and SPARC in those samples (**Figure 8**). In each case, the correlation was significant according to the p and R values.

**Figure 6.**
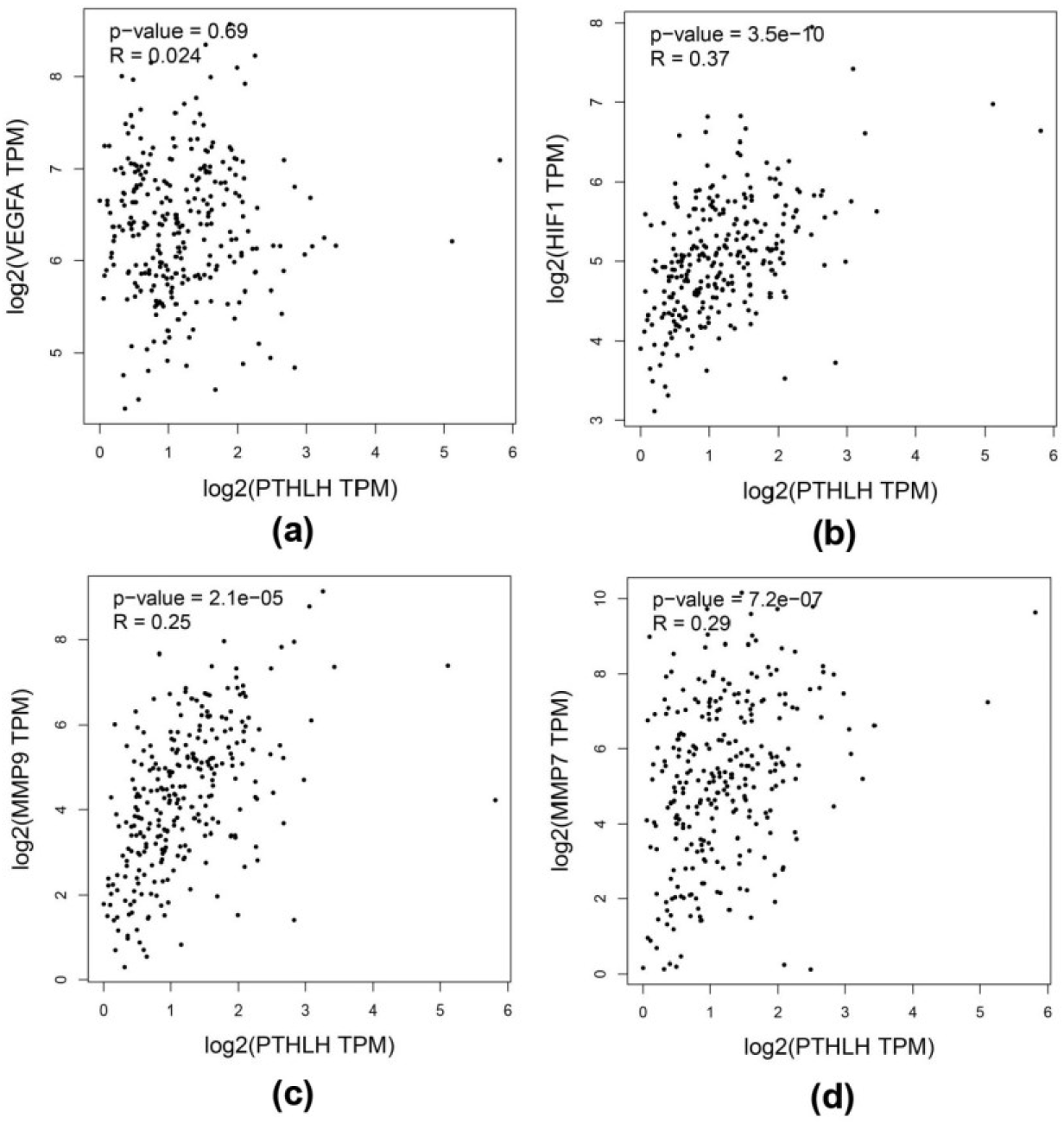
Correlation between the PTHLH gene and markers associated with angiogenesis. The graphs show the correlation between PTHLH expression with **(a)** VEGFA, **(b)** HIF-1, **(c)** MMP-9, and **(d)** MMP-7 using GEPIA2 platform that analyzes information from CRC tumor samples from the TCGA repository.

**Figure 7.**
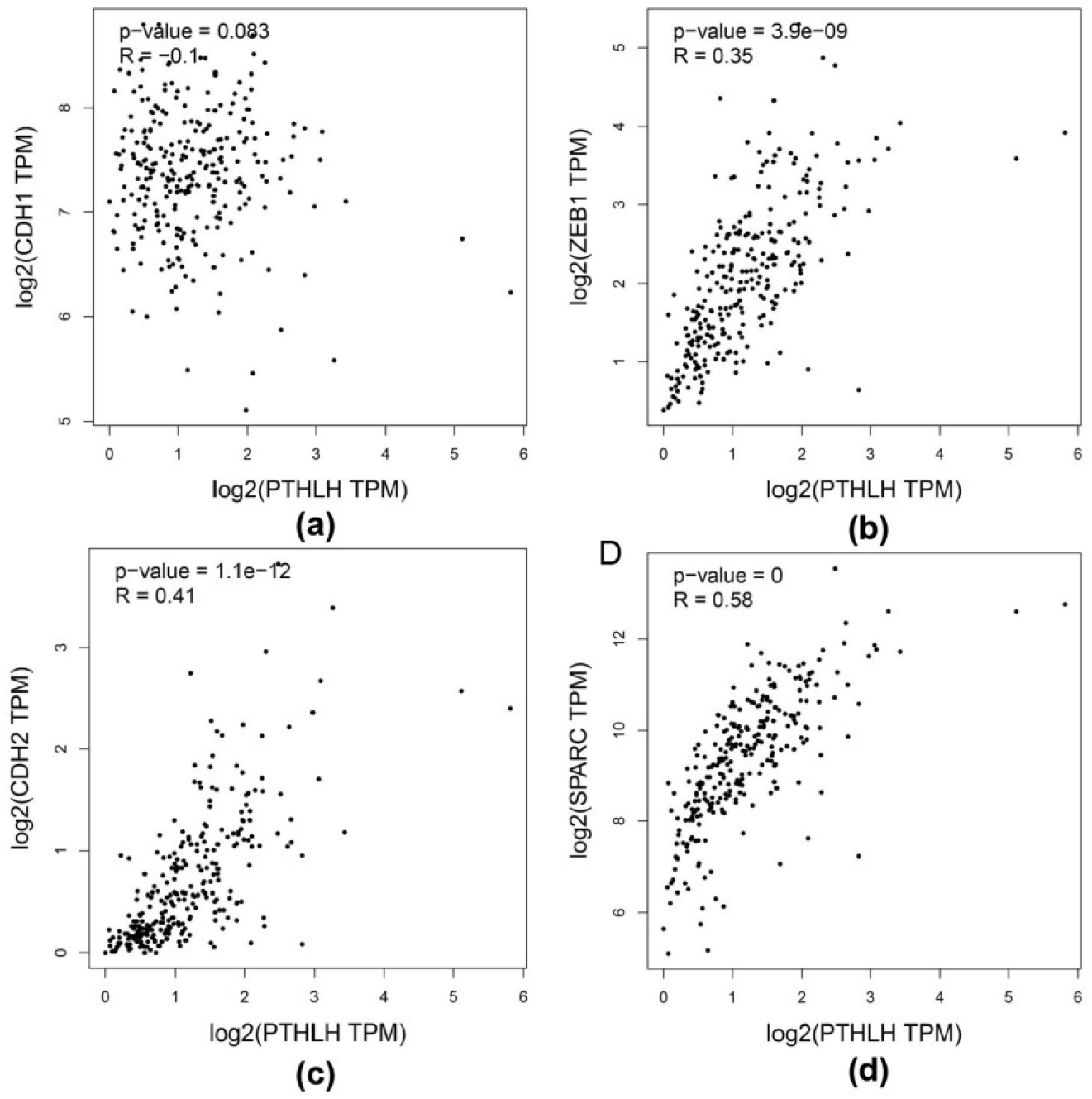
Correlation between the PTHLH gene and markers associated with EMT. The graphs show the correlation of PTHLH gene expression with **(a)** the E-cadherin gene (CDH1), **(b)** the ZEB1 gene, **(c)** the N-cadherin gene (CDH2), and **(d)** the SPARC gene, using the GEPIA2 platform that analyzes data from CRC tumor samples from the TCGA repository.

**Figure 8.**
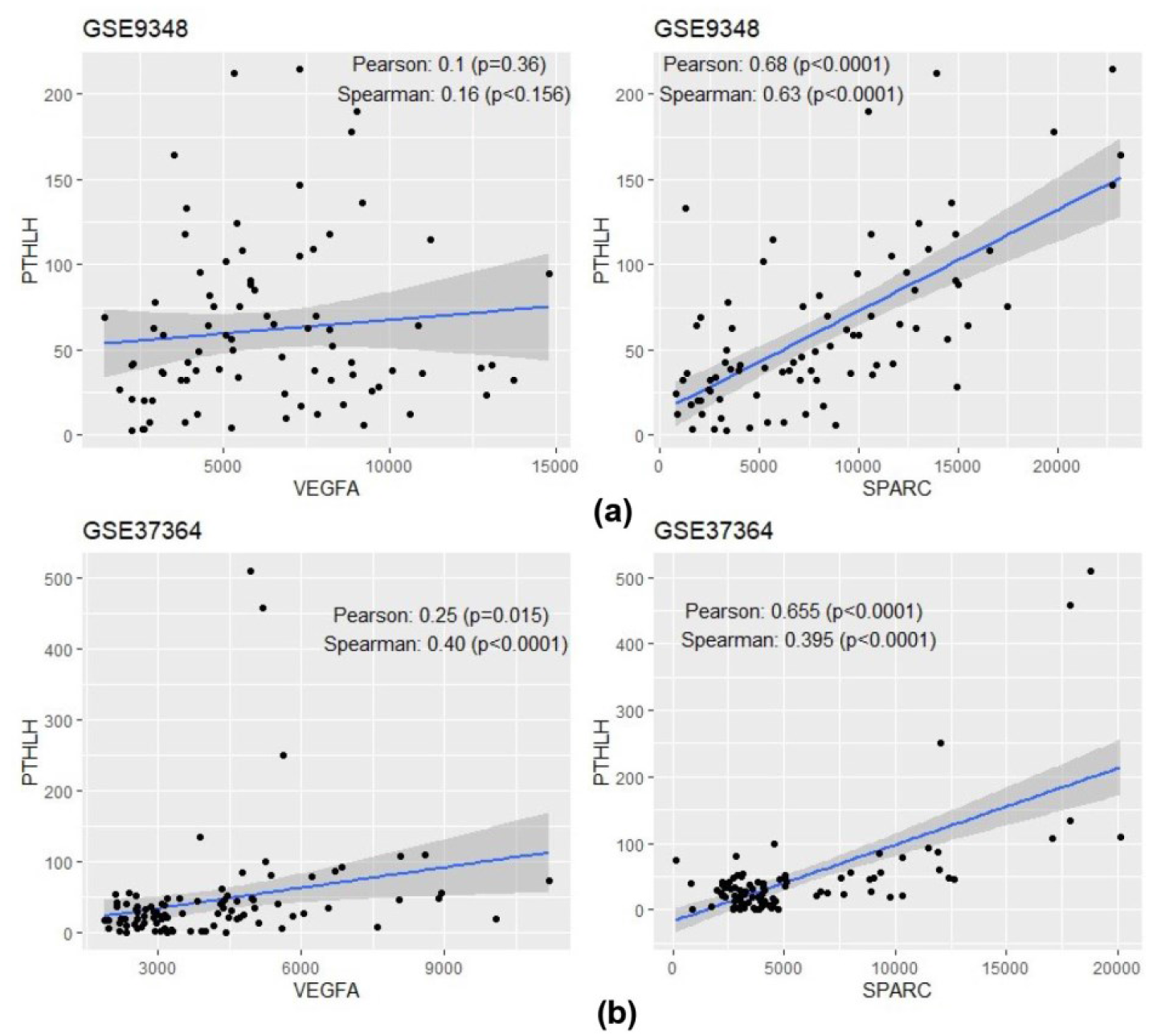
PTHLH correlates with VEGFA and SPARC in CRC clinical samples from the accessions queried. Analysis of GEO databases in which PTHLH is upregulated shows that PTHLH correlates positively with **(a)** SPARC (GSE9348) and with **(b)** VEGFA and SPARC (GSE37364) in CRC tumor samples.

### 3.3 Network biology study of the interaction between PTHrP and markers of angiogenesis and EMT

Based on our previous findings that demonstrate that PTHrP increases the expression of SPARC in HCT116 and HMEC-1 cell lines derived from CRC and human microvasculature, respectively. [11], a network biology approach was used to further study the relationship between PTHrP and SPARC and to visualize the interaction of proteins related to those. For this reason, the possibility of a regulatory network between both factors of the tumor microenvironment was inferred. Cytoscape is an open-source program used for the exploration (visualization, analysis, and interpretation) of biological and biomedical networks that allows the analysis of data from different sources in the form of interconnections supported by experimental and bibliographic associations, allowing the prediction of the biological function of biomarkers and their regulatory pathways [29]. To integrate the STRING 11.5 database and the Cytoscape network analysis platform, the stringApp application was downloaded, a Cytoscape plugin that facilitates the import of STRING networks and the integration with additional data provided by the user in a single workflow. To elucidate the possible regulatory network between PTHLH and SPARC, the interaction between both genes and the different proteins associated with these factors in the aforementioned databases was explored through StringApp [19].

As seen in **Figure 9a**, the search for proteins associated with PTHLH and SPARC generated a network of 650 interactions. The proteins that presented the highest degree of interaction between PTHLH and SPARC were then selected and a smaller network was created, in which VEGFA is visualized as a node with a high interaction score between both factors (**Figure 9b**).

**Figure 9.**
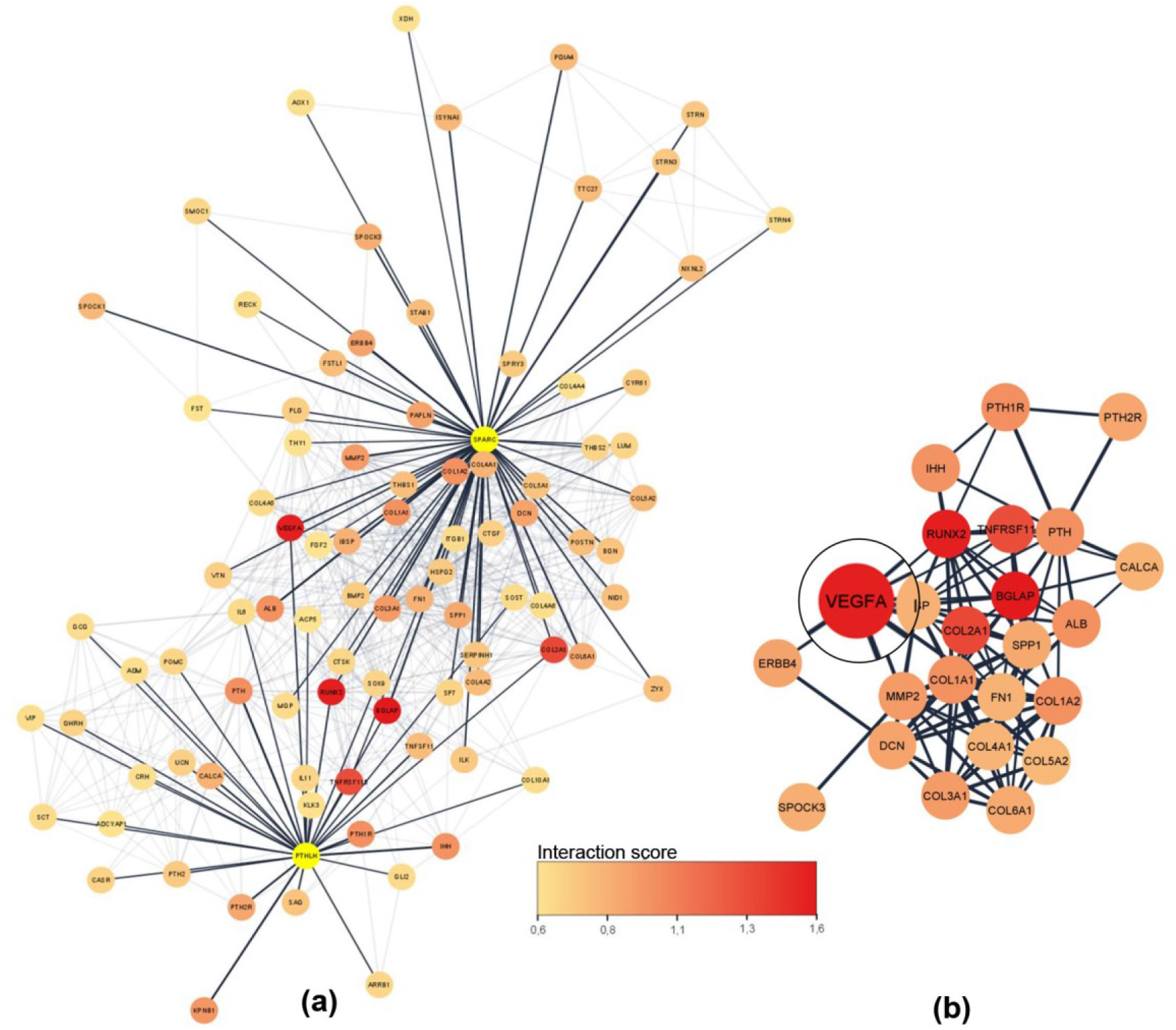
Interaction network between PTHLH and SPARC. STRING *protein search* was used as the search source, with the input “PTHrP SPARC”. The network was created with a confidence score cutoff greater than 0.6 and a maximum of 100 proteins (nodes) establishing a network of 92 nodes and 650 interactions. **(a)** PTHLH and SPARC (yellow) are shown with their direct interacting proteins; **(b)** Network formed with the genes with the highest interaction score (IS) of the “PTHrP SPARC” network (IS> 0.95).

Finally, the implication of the previously obtained relevant DEGs in CRC was established with the group of genes with the highest interaction in the PTHLH and SPARC network. Based on the results of the functional enrichment that showed the association of the genes positively regulated in CRC in the four data sets with the processes of angiogenesis and EMT, the link between these genes and the genes associated with PTHLH and SPARC was explored.

As can be seen in **Figure 10**, an interaction network of the positively regulated genes was created with a confidence score limit greater than 0.4 and a maximum of 100 proteins (nodes), which generated a network of 789 interactions. Using the CytoNCA application for centrality analysis shows VEGF as a central node in the network of these genes in the clinical samples evaluated.

**Figure 10.**
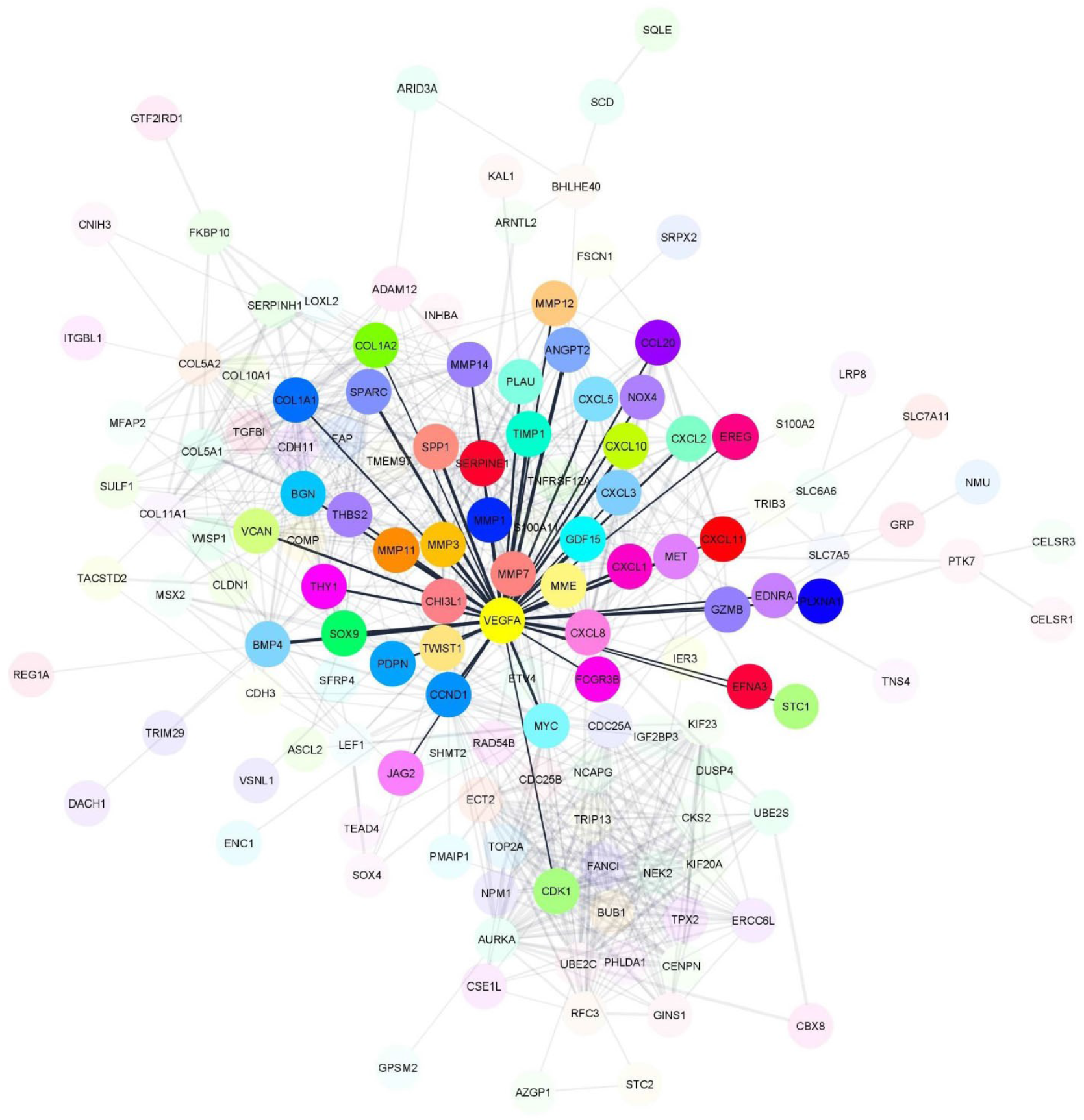
The network obtained from the positively regulated genes in the evaluated clinical samples of CRC shows VEGFA as a central node. CytoNCA app was used to find the genes with the highest network centrality score.

Clusters of nodes from this network that are highly interconnected were then selected using the Cytoscape MCODE plugin [20]. The identification of densely connected nodes in a network is useful for identifying biological modules, such as complexes or signaling pathways. In this case, MCODE identified 3 functionally homogeneous clusters from the interaction network of positively regulated genes in CRC (**Figure 11**). The visualization of the network obtained from the proteins most associated with PTHLH and SPARC (see **Figure 9**), and the network of the positively regulated genes in the four CRC data sets, shows VEGFA as the shared node between both networks. Furthermore, the MCODE clustering of the CRC network shows the existence of a homogeneous cluster (Cluster 2) containing SPARC and VEGFA as part of the same functional module (**Figure 11b**) associated with EMT and angiogenesis (**Figure 12**). This linkage includes experimental/biochemical data, association in selected databases and joint mention in Pubmed abstracts.

**Figure 11.**
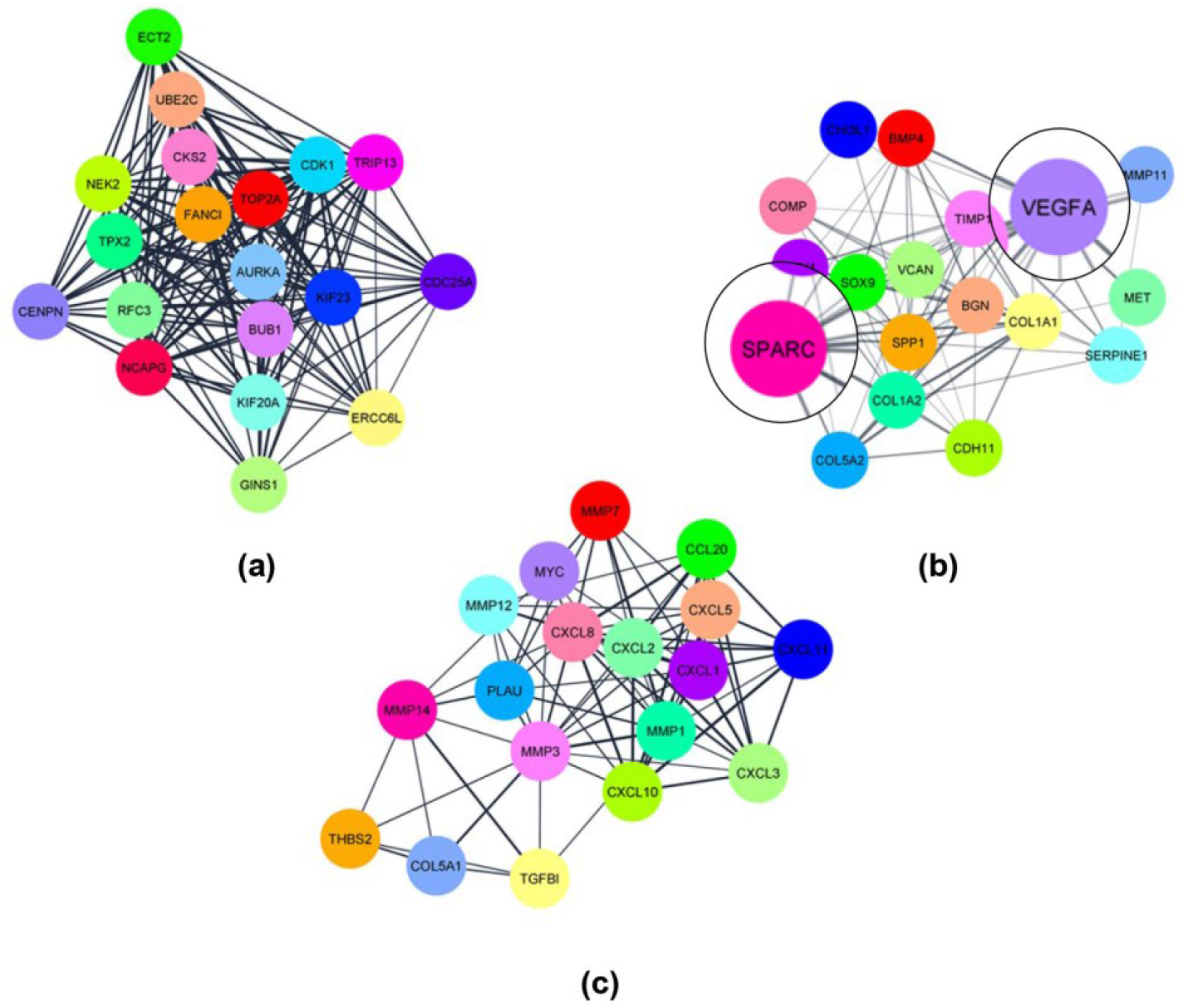
Clustering analysis of the network of positively regulated genes in CRC patients shows that VEGF and SPARC are in the same functional module. MCODE was used to obtain the 3 most homogeneous clusters in the network. **(a)** Module 1 with a score of 17.889 (19 nodes, 161 edges); **(b)** Module 2 with a score of 10.471 (18 nodes, 89 edges); **(c)** Module 3 with a score of 9.647 (18 nodes, 82 edges). VEGFA and SPARC are part of the same functional module (Module 2). The parameters used were cut-off degree: 2, node density cut-off: 0.1; node score cut-off: 0.2; K-core: 2.

**Figure 12.**
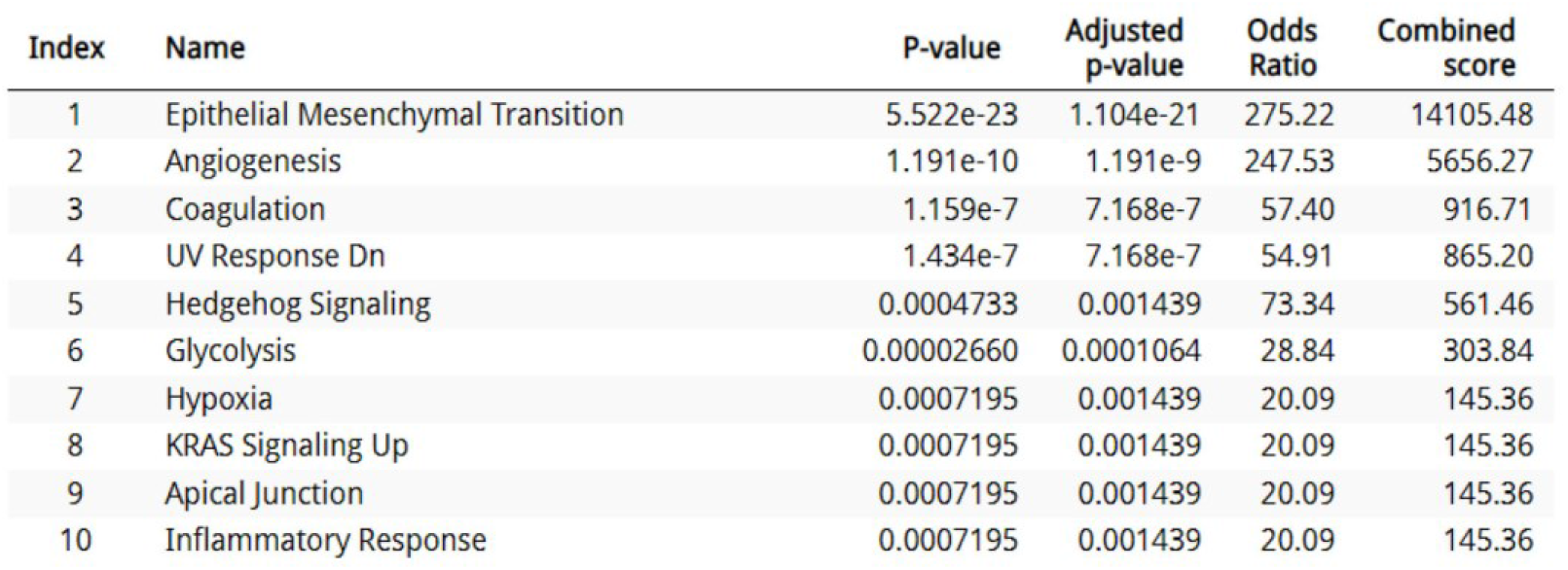
MSigDB Hallmark 2020 molecular signature database shows the 10 most significant terms associated with genes in Module 2 obtained by MCODE. Module 2 is associated with pathways involved in EMT and angiogenesis. The p-values and q-values represent the probability of association and the Bejamini-Hochberg adjusted probability, respectively.

### 3.4 Prognostic significance of PTHLH, SPARC, and VEGFA expression in colorectal cancer

To assess the clinical relevance of these findings Survival analysis was performed to evaluate the prognostic significance of PTHLH, SPARC, and VEGFA expression in CRC patients. High expression levels of each individual gene were significantly associated with reduced overall survival. Specifically, increased PTHLH expression was associated with poorer survival outcomes (HR = 1.37, 95% CI: 1.11–1.68, p = 0.0028). Similarly, elevated SPARC expression correlated with decreased overall survival (HR = 1.34, 95% CI: 1.09–1.65, p = 0.0048), as did VEGFA expression (HR = 1.35, 95% CI: 1.10–1.66, p = 0.0039). To further investigate the combined prognostic value of these genes, a multigene signature was constructed using the mean expression of PTHLH, SPARC, and VEGFA. This combined signature demonstrated a markedly stronger association with overall survival compared to each gene individually (HR = 1.66, 95% CI: 1.35–2.04, p = 1.2 × 10^−6^) (**Figure 13**). These results indicate that while each gene individually exhibits a moderate prognostic effect, their combined expression provides enhanced predictive power, supporting a cooperative role of these factors in colorectal cancer progression.

**Figure 13.**
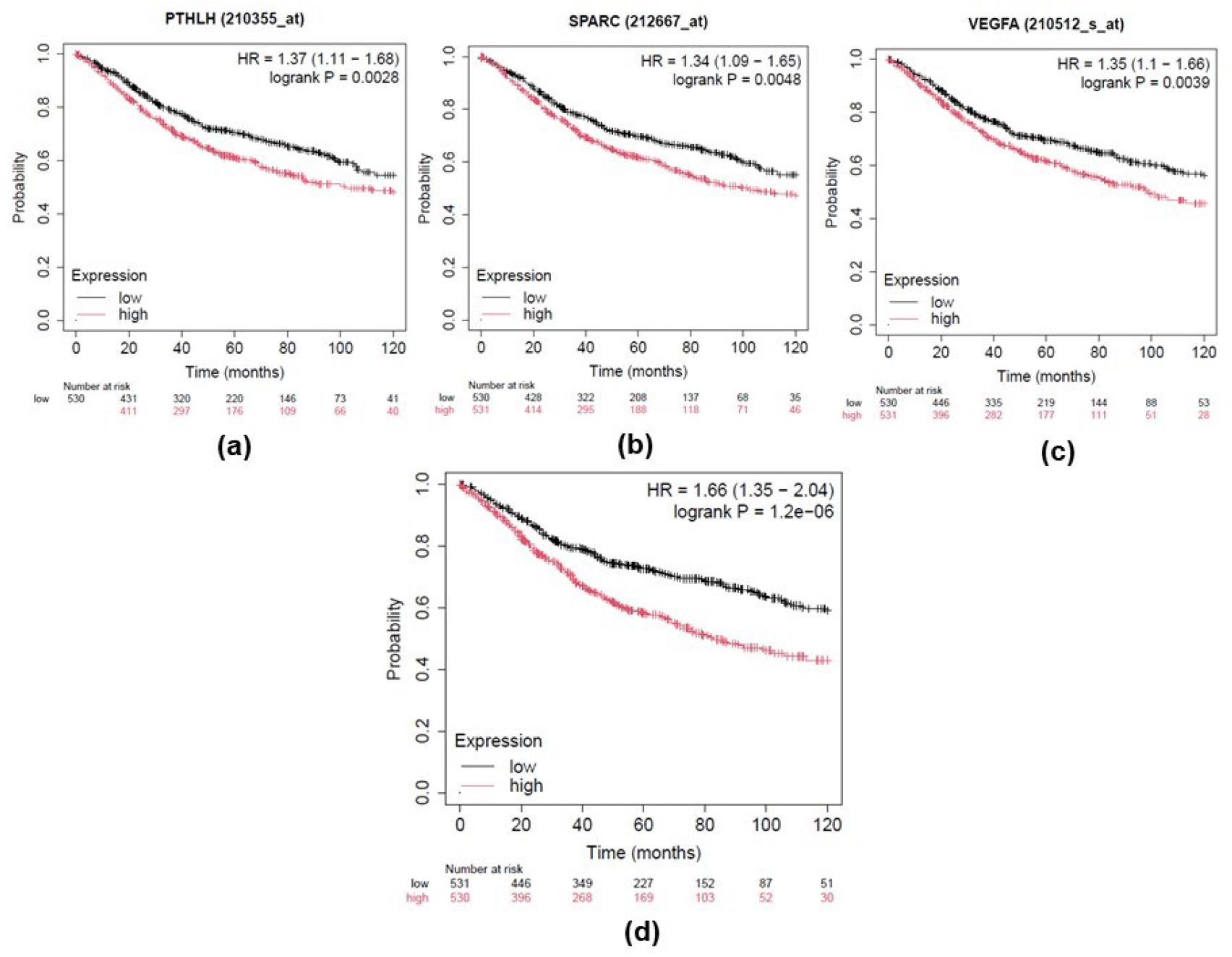
Kaplan-Meier survival analysis of PTHLH, SPARC, and VEGFA expression in CRC patients. Kaplan-Meier curves showing overall survival (OS) according to gene expression levels. Patients were stratified into high- and low-expression groups using the auto-selected best cutoff. **(a)** High expression of PTHLH was significantly associated with reduced overall survival (HR = 1.37, 95% CI: 1.11–1.68, p = 0.0028). **(b)** Elevated SPARC expression was associated with poorer survival outcomes (HR = 1.34, 95% CI: 1.09–1.65, p = 0.0048). **(c)** Increased VEGFA expression correlated with decreased overall survival (HR = 1.35, 95% CI: 1.10-1.66, p = 0.0039). **(d)** Combined analysis of PTHLH, SPARC, and VEGFA expression demonstrated a stronger prognostic effect compared to individual genes (HR = 1.66, 95% CI: 1.35–2.04, p = 1.2 × 1^−6^0).

### 4. Discussion

Since the emergence of the TCGA project, which includes descriptive and comparative studies of clinical samples from 33 types of cancer, the number of available online platforms and data analysis programs has grown in parallel, condensing the enormous amount of information published in recent years. Other authors and we have conducted exploratory studies for the detection of biomarkers in CRC with a workflow similar to that applied in this work [30-33], however, few studies have used multiple microarray data sets to study hub genes and there are no studies that have addressed the relationship of PTHrP with relevant CRC genes, analyzing human samples.

In this study, we performed an integrative analysis of multiple transcriptomic datasets to investigate the potential role of PTHLH in CRC. Our results identified a set of differentially expressed genes consistently shared across datasets and revealed that PTHLH expression is associated with molecular pathways related to extracellular matrix remodeling, angiogenesis, and tumor progression. The results of the analysis of the evaluated data from CRC patient samples reveal that 176 positively regulated genes and 254 downregulated genes are shared by the four sets. Although the genes shared by the four data sets were used for the analysis, it is noteworthy to consider that in the GSE37364 data set and in the GSE9348 data set, PTHLH is positively regulated (Log_2_FC=2.34; adjusted p <0.05 and Log_2_FC=1.52; adjusted p <0.05 respectively). For this analysis, the samples were not classified based on other parameters such as tumor stage, occurrence of metastasis, and adjuvant treatments before sampling, among others. For this reason, it is necessary to further investigate whether PTHrP positive regulation is influenced by these patients’ features. In addition, unlike the GSE41258 and GSE68468, the GSE37364 and GSE9348 platforms belong to the Affymetrix Human Genome U133 Plus 2.0 Array model, which has a greater number of search probes than the Affymetrix Human Genome U133A Array platform. The fact that we select four data sets from independent repositories, could explain the absence of information on PTHLH in the two data sets mentioned.

Functional enrichment and network analyses suggested that positively regulated genes in CRC samples are linked to biological processes previously associated with tumor aggressiveness. In this context, PTHLH was identified as an EMT-associated gene and showed significant associations with angiogenesis-related pathways in colorectal cancer. These findings are consistent with prior *in vitro* and *in vivo* evidence indicating that PTHrP may contribute to tumor progression through modulation of the tumor microenvironment.

Further analysis using independent datasets revealed a positive correlation between PTHLH and SPARC expression, supporting a potential association between these factors in CRC. Although correlations with metalloproteases such as MMP-7 and MMP-9 were statistically significant, their relatively low correlation coefficients suggest that these relationships should be interpreted with caution. Similarly, while no consistent correlation between PTHLH and VEGFA was observed across all datasets, alternative analyses and network-based approaches indicated that VEGFA may still be functionally connected within the same regulatory context. These discrepancies may reflect differences in dataset composition or analytical methodologies.

As previously mentioned, SPARC is closely related to the extracellular matrix remodeling, invasion, and migration of tumor cells and vascular development in the TME [34]. Several investigations associate SPARC and VEGF expression. In colon cancer, it was observed that SPARC de-repress the expression of VEGF modulating angiogenesis [34]. Although SPARC showed an angiogenic effect, other authors observed that this peptide inhibits the activity of VEGF on endothelial cells influencing the angiogenic process [35]. Chandrasekaran and colleagues observed that SPARC interacts with the binding site of VEGF in the VEGFR-1 (Vascular endothelial growth factor receptor Type 1) inhibiting the proliferation of endothelial cells [35]. It was also demonstrated that SPARC activity in some circumstances promotes angiogenesis [36]. In fact, in ovarian carcinoma, an overexpression of SPARC and VEGF was detected [37]. Concerning this, Kato and collaborators suggested a dual role of SPARC on VEGF functions [34]. However, to date the information regarding the interrelation of both factors in CRC remains unclear, and this work complements and reinforces the results previously obtained by our research group and other groups on this topic [10,11,33,39].

Importantly, the clinical relevance of these observations was supported by survival analysis. High expression levels of PTHLH, SPARC, and VEGFA were each significantly associated with reduced overall survival, although with moderate effect sizes. Notably, the combined gene signature demonstrated a substantially stronger prognostic impact compared to individual genes, indicating that their joint expression provides enhanced predictive value. These results support a cooperative role of these factors and highlight the potential of multigene signatures to better capture the complexity of tumor biology in CRC. Despite these findings, it is important to recognize that the analyses were based on publicly available transcriptomic datasets, which may introduce variability due to differences in experimental platforms, sample processing, and patient characteristics. Furthermore, the present study is based on associative analyses and does not establish causal relationships. Therefore, further experimental validation will be necessary to elucidate the mechanistic role of PTHLH and its interaction with SPARC and VEGFA.

### 5. Conclusion

Our results suggest that PTHLH expression is associated with molecular pathways involved in EMT and angiogenesis in CRC. Functional enrichment analyses and network-based approaches indicate that SPARC and VEGFA are part of a closely connected functional group, with VEGFA emerging as a highly interacting node within the PTHLH-associated network. These findings support the existence of a coordinated PTHrP-SPARC-VEGF axis that may contribute to tumor progression. Importantly, the combined expression of these genes demonstrates significant prognostic value, highlighting this molecular signature as a potential candidate for further investigation in colorectal cancer.

## Author contributions

Conceptualization, PC, CG; methodology, PC; software, PC; validation, PC and MB ND; formal analysis, PC; investigation, PC, MB ND; resources, CG; data curation, PC; writing-original draft preparation, PC, MB ND, CB and CG; writing-review and editing, CG; visualization, PC; supervision, CG; project administration, CG; funding acquisition, CG. All authors have read and agreed to the published version of the manuscript.

## Acknowledgments

The authors thank Diego Nabaes Jodar for advice on the statistical analysis of the manuscript.

## Ethical approval

The clinical samples selected for this study are located in public repositories and have the corresponding ethical endorsement of the article where they were published.

